# Was facial width-to-height ratio subject to sexual selection pressures? A life course approach

**DOI:** 10.1101/2020.09.24.311324

**Authors:** Carolyn R. Hodges-Simeon, Graham Albert, George B. Richardson, Timothy S. McHale, Seth M. Weinberg, Michael Gurven, Steven J.C. Gaulin

**Author notes:** Corresponding author (CH). C.R.H. and S.M.W. collected the data. S.M.W., M.G., and S.J.C.G. were involved in planning and supervised the work. C.R.H. and G.A. organized the data, performed the analysis, drafted the manuscript and designed the figures. C.R.H, G.A., G.B.R., T.S.M., S.M.W., M.G., and S.J.C.G. aided in interpreting the results and worked on the manuscript.

## Abstract

Sexual selection researchers have traditionally focused on adult sex differences; however, the schedule and pattern of sex-specific ontogeny can provide insights unobtainable from an exclusive focus on adults. Recently, it has been debated whether facial width-to-height ratio (fWHR; bi-zygomatic breadth divided by midface height) is a human secondary sexual characteristic (SSC). Here, we review current evidence, then address this debate using ontogenetic evidence, which has been under-explored in fWHR research. Facial measurements collected from males and females aged 3 to 40 (Study 1; US, *n=*2449), and 7 to 21 (Study 2; Bolivia, *n*=179) were used to calculate three fWHR variants (which we call fWHR*nasion*, fWHR*stomion*, and fWHR*brow*) and two other common facial masculinity ratios (facial width-to-lower-face-height ratio, fWHR*lower,* and cheekbone prominence). We test whether the observed pattern of facial development exhibits patterns indicative of SSCs, i.e. differential adolescent growth in either male or female facial morphology leading to an adult sex difference. Results showed that only fWHR*lower* exhibited both adult sex differences as well as the classic pattern of ontogeny for SSCs—greater lower-face growth in male adolescents relative to females. fWHR*brow* was significantly wider among both pre- and post-pubertal males in the 2D sample; post-hoc analyses revealed that the effect was driven by large sex differences in brow height, with females having higher placed brows than males across ages. In both samples, all fWHR measures were inversely associated with age; that is, human facial growth is characterized by greater relative growth in the mid-face and lower face relative to facial width. This trend continues even into middle adulthood. BMI was also a positive predictor of most of the ratios across ages, with greater BMI associated with wider faces. Researchers collecting data on fWHR should target fWHR*lower* and fWHR*brow* and should control for both age and BMI.

## Introduction

Charles Darwin (1872) used the term *secondary sexual characteristic* (SSC) to refer to traits that evolve by sexual selection, and which contribute to an individual’s reproductive success through deterring competitors (i.e., intrasexual selection; Andersson, 1994; Buss 1988; Lindenfors & Tullberg, 2011; Puts, 2010) or attracting mates (i.e., intersexual selection; Andersson, 1994; Buss, 1989). Sexual selection is the primary explanatory framework for the evolution of sex differences across species, including humans (e.g., Andersson, 1994; Conroy-Beam et al., 2015; Lassek & Gaulin, 2009; Penton-Voak et al., 1999; Plavcan, 2012; Puts et al., 2007; Puts, 2010).

In 2007, Weston et al. proposed a new human SSC—facial width-to-height ratio (fWHR), or the width of the face (between the left and right zygion) divided by the length of the mid-face (from the nasion to the prosthion, referred to as fWHR*nasion* in the current analyses; see Table 1 and Fig 1 for measurement variants) based on identification of sex differences in a sample of South African crania. Since then, this and similar facial metrics have gained increasing attention^1^ in psychology, biological anthropology, and other fields for its persistent association with an array of behavioral, psychosocial, and anatomical traits (e.g., Carré, & McCormick, 2008; Carré, McCormick, & Mondloch 2009; Gómez-Valdés et al., 2013; Hodges-Simeon, Sobraske, Samore, Gurven, & Gaulin, 2016). A number of recent studies, however, highlight inconsistencies in the findings (Lefevre et al., 2012; Kosinski, 2017; Özener, 2012) and it is now currently debated whether fWHR should be characterized as a SSC (Dixson, 2018; Hodges-Simeon et al., 2016; Hodges-Simeon et al., 2018; Kramer, 2017; Welker et al., 2016). We review the current debate, and then argue that important insights may be gained from an ontogenetic approach, which should inform any conclusions drawn from adult populations.

### Is fWHR a secondary sexual characteristic (SSC)?

Evolutionary biologists emphasize three joint criteria to assess whether a trait is a product of sexual selection rather than an alternative process (e.g., genetic drift, pleiotropic byproduct; Järvi et al., 1987).

1. SSCs should be sexually dimorphic, at least during the period(s) of mating competition (Andersson, 1994). Weston et al. (2007) first described sex differences in dry bone fWHR among a sample of native southern African crania. However, since then, identification of adult sex differences in fWHR have been inconsistent; several studies have found significant sex differences (Carré, & McCormick, 2008; Weston et al., 2007), while others have not (Gómez-Valdés et al., 2013; Kramer, Jones, & Ward, 2012; Kramer, 2015; Kramer, 2017; Ozener, 2012; Robertson, Kingsley, & Ford, 2017). A recent meta-analysis of these findings indicated a significant adult sex difference in fWHR, but the magnitude of the effect was small (*mean weighted effect size* = 0.11; Geniole et al., 2015). For comparison, three traits that likely are SSCs—stature, voice pitch, and muscularity—show much larger sex differences, with effect sizes of 1.63 (height, across 53 nations; Lippa, 2009), 2.38 (vocal fundamental frequency; Vogel et al., 2009), and 2.5 (arm muscle volume; Lassek and Gaulin, 2009).

2. SSCs should increase success in mating competition, leading to higher reproductive success (or proxies thereof, such as mating success or judgments of attractiveness; Apicella, Marlowe, & Feinberg, 2007; Hughes, Dispenza, Gallup, 2004; Gontard-Danek & Møller, 1999). The evidence that men with greater fWHRs have greater reproductive success has been mixed. Studies have shown that men with greater fWHR have greater mating success (Valentine, Li, Penke, & Perrett, 2014), increased sex drive (Arnocky et al., 2017), and more children (Loehr & O’hara, 2013); whereas other studies have not identified a relationship between men’s fWHR and number of children (Gómez-Valdés et al., 2013).

Weston et al. (2007) originally proposed that a larger fWHR in males (i.e., wider face relative to midface height) may have evolved by intersexual selection (i.e., female choice); however, a meta-analysis showed a significant *negative* relationship between fWHR and physical attractiveness ratings across 8 studies; i.e., women judged men with wider faces to be *less* attractive (Geniole et al., 2015). In contrast, there is more compelling support for the notion that fWHR was shaped by intrasexual competition among males. Wider faces seem to be reliably associated with a suite of behavioral traits involved in physical competition (e.g., aggressive behavior in sports; Carré, & McCormick, 2008; Carré, McCormick, & Mondloch 2009) and aggression in both naturalistic and laboratory settings (Carré, & McCormick, 2008; Geniole et al., 2015; Welker, Goetz, Galicia, Liphardt, & Carré, 2015; Zilioli et al., 2015). While several studies found no relationship between fWHR and aggression-linked traits (e.g., Lewis et al., 2012), self-reported aggression (Özener, 2012), or behavioral measures of aggression (Deanor et al, 2012), meta-analyses show a strong and consistent relationship between higher male fWHR and *perceptions* of aggressiveness, fighting ability, masculinity, dominance, and threat by both male and female raters (*r* = .13-.46; Geniole et al., 2015; Geniole & Mccormick 2015; Zilioli et al., 2015). In addition, fWHR is linked to measures of dominance, status, or assertiveness among capuchin monkeys (*Sapajus spp:* Lefevre et al., 2014), macaques (*Macaca mulatta*; Altschul et al., 2019; Borgi & Majalo, 2016), and bonobos (*Pan paniscus*; Martin et al., 2019).

3. SSCs often co-occur with a suite of other behavioral, physiological, and morphological traits that jointly contribute to a particular mating strategy (Geniole et al., 2015). For instance, selection on larger body size and muscle mass in males (relative to females) usually co-occurs with the behavioral inclination to use these weapons (Sell et al., 2009; Sell et al., 2016), yet fWHR was not associated with grip strength in either sex in a recent study (MacDonell et al., 2018). Some research suggests fWHR is best understood as a predictor of behavioral strategies that promote status-seeking (Lewis et al., 2012), power, and resource acquisition, such as willingness to cheat or exploit the trust of others to increase financial gain (Geniole, Keyes, Carré & McCormick, 2014; Haselhuhn & Wong, 2011; Jia et al., 2014; Stirrat & Perrett, 2010), risk-taking (Welker et al., 2015), and narcissism (Noser et al., 2018; however, see Kosinski, 2017). Many authors reason that the link between these behavioral strategies and fWHR stems from their joint regulation by testosterone (Bird et al., 2016; Carré & McCormick, 2008). However, amongst adult males, a meta-analysis showed no significant relationship between fWHR and basal T concentrations (Bird et al., 2016) or androgen receptor gene polymorphisms (Eisenbruch et al., 2017). For reactive T (i.e., change in T in response to challenge), Lefevre et al. (2013) found a positive association with fWHR, yet Bird et al. (2016) and Kordsmeyer et al. (2019) did not. Research on wider face shape and higher prenatal testosterone is promising (Bulygina et al., 2006; Weinberg et al., 2014; Whitehouse et al., 2016), but further studies on hormonal correlates fWHR are needed.

In summary, for each of the three criteria useful in identifying SSCs, the previously published evidence is weak, conflicting or ambiguous. The first criterion has been under-examined in the literature; that is, the majority of studies focus on adult sex differences. In the present study, we examine the developmental pattern of fWHR (as well as several other facial masculinity ratios) to assess whether these ratios demonstrate sex-specific changes that occur in tandem with the commencement of sexual maturation.

### Ontogenetic perspectives on sexual selection

Evolution and ontogeny are closely intertwined because intra- and interspecific evolutionary change in the adult phenotype occurs by means of changing schedules of ontogeny (Bogin, 1999; Gould, 1977; Leigh, 1995). For example, sex differences in adult height can be explained quantitatively by the delayed onset, increased rate, and longer duration of the adolescent growth spurt in males compared with females (Hauspie & Roelants, 2012). This sex-specific pattern of growth suggests that selection for a later and longer growth spurt in males outweighed the costs of later reproduction. Research on fWHR—as well as on sexual selection more generally—has almost exclusively drawn from studies of adult males and females; however, the schedule and pattern of sex-specific development can provide insights on sexual selection pressures unobtainable from studies limited to adults (e.g., Badyaev, 2002; Mank et al., 2010; Taylor, 1997; Hodges-Simeon et al. 2014, 2015). Several types of ontogenetic data should be particularly useful to those interested in sexual selection pressures.

First, SSCs should develop in temporal contiguity with the commencement of mating competition. For some species, this may occur during defined mating seasons (e.g., Burger et al., 2013; Galea et al., 1994; Järvi et al., 1987; Pyter et al., 2005; Smith et al., 1997) or transient exposure to potential mates (e.g., Amstislavskaya & Popova, 2004; Roney et al., 2007), while in others SSC development may canalize during reproductive maturation (i.e., puberty in humans; Hochberg, 2012; Hodges-Simeon et al. 2013). Thus far, only Weston et al. (2007) has examined sex differences in fWHR prior to adulthood (although see Kesterke et al., 2016; Koudelová et al., 2019; and Matthews et al., 2018 for sex-specific development in non-ratio facial dimensions); therefore, our primary goal is to determine if fWHR (along with several other commonly used facial masculinity ratios) exhibits sex-specific divergence during puberty. To further clarify the developmental pattern and shed light on the role of sexual selection, we assess whether sex differences, if present, arise from male-specific or female-specific growth as a proxy for selection pressures acting on males versus females.

Second, male-specific trait development during or before mating competition is orchestrated by androgens such as testosterone (e.g., Galea et al., 1999; Hodges-Simeon, Gurven, & Gaulin, 2015; Marečková et al., 2011; Marečková et al., 2015; Pyter et al., 2006; Spritzer & Galea, 2007; Verdonck et al., 1999); thus, an association between testosterone and trait development of masculine features is often treated as evidence for sexual selection in mammalian males (Folstad & Karter, 1992; Bird et al., 2016). Few studies, however, have examined the association between fWHR and testosterone prior to adulthood. Hodges-Simeon et al. (2016) showed that among adolescents, fWHR was not associated with age, and only weakly with testosterone (see also Welker et al., 2016; Hodges-Simeon et al., 2018). This is in stark contrast to more established SSCs (e.g., voice pitch, muscle mass), which show very strong associations with testosterone and age during the adolescent period—a phase when testosterone increases by an order of magnitude in only 5 to 9 years (Butler et al., 1989; Elmlinger et al., 2004; Kelsey et al., 2014; Hodges-Simeon et al., 2015).

Third, if fWHR is a SSC, then it should exhibit ontogenetic patterns similar to other human SSCs. SSCs typically emerge together during puberty because they form a functional suite of tactics supporting success in mating competition. Thus, we should see males’ and females’ fWHR diverge in the phase between puberty and adulthood—i.e., adolescence (or potentially in the period between adrenarche and puberty, called juvenility or middle childhood; Bogin, 1999; Pereira & Fairbanks, 1993). The pattern of development in males may also exhibit a “spurt” (i.e., a period of increasing growth velocity), which is descriptive of the growth pattern of male muscle mass, height, and voice pitch (Hodges-Simeon et al., 2016). This pattern is likely due to regulation by testosterone, which itself shows a pronounced spurt (Hodges-Simeon et al., 2016). Currently, there is a deficit of findings on the ontogeny of fWHR and other commonly used facial masculinity ratios, which this research seeks to address.

### Aims and predictions of the present research

We propose four aims and associated predictions for the present study. Our first goal is to test for the presence or absence of adult sexual dimorphism in fWHR in a large, homogenous (i.e., European-Caucasian; *N* = 1,477, aged 22-40) sample. Previous studies have diverged, with some showing a significant sex difference (Carré, & McCormick, 2008, *N* = 88; Weston et al., 2007, *N* = 121) and others not (Gómez-Valdés et al., 2013, *N* = 4,960; Kramer, Jones, & Ward, 2012, *N* = 415; Kramer, 2015, *N* = 3,481; Kramer, 2017, *N* = 7,941; Ozener, 2012, *N* = 470; Robertson, Kingsley, & Ford, 2017, *N* = 444), which utilize 2D, 3D, and dry bone skull samples. Kramer et al. (2017) has targeted the largest sample of fWHR in dry bone skulls thus far (*N* = 7,941), showing small but significant sex differences in fWHR in East Asian but not any other populations. We offer the largest sample size to date for fWHR from soft tissue, three-dimensional faces. This is an important complement to the literature on dry bone morphology, as sexual dimorphism may stem not only from divergence in craniofacial growth, but also sex-specific patterns of muscle and fat deposition (Lassek & Gaulin, 2009; Woods & Wong, 2016).

Our second aim is to examine sex differences and sex-specific growth in fWHR in sub-adult age groups (i.e., childhood, juvenility, and adolescence), and to determine if sex differences in fWHR are due to male-specific or female-specific growth—questions that have not yet been addressed in the literature. For most human SSCs, pre-pubertal groups show little-to-no difference, while those in later adolescence and adulthood exhibit more observable differences. Sex differences may derive from male-specific growth (i.e., male features growing faster or longer than females’), female-specific growth (i.e., female features growing faster or longer than males’), or a combination of the two. To this end, we measure fWHR among sub-adult males and females in two populations: the large European-Caucasian sample of 3D facial scans (ages 3 to 21) and an indigenous Bolivian Tsimane sample of 2D front-facing photographs (ages 7 to 21).

Our third goal is to examine variation in fWHR growth velocity (i.e., acceleration) across ages as the pattern of ontogeny may yield additional insight. In particular, human male SSCs typically show evidence of a growth spurt during adolescence—rapid acceleration followed by deceleration—due to the influence of testosterone on this trait. This was previously examined in our Tsimane dataset (Hodges-Simeon et al., 2016), which showed no evidence of a growth spurt in several different fWHR ratios. However, because this sample was small, we address the question again here in our 3D dataset, which offers a larger N.

Our fourth goal is to examine sex differences and sex-specific development in several other commonly used facial masculinity ratios that, unlike fWHR, incorporate mandibular proportions (Lefevre et al., 2012; Penton-Voak et al., 2001; Little et al., 2016): the ratio of bizygomatic facial width to the width of the face at the mouth (“cheekbone prominence”) and the ratio of bizygomatic width to morphological face height (nasion to bottom of chin; “fWHR*lower*”, see Fig 1). fWHR*lower* and cheekbone prominence are smaller in adult men compared to women (Lefevre et al., 2012) because of the relatively larger size of the male mandible. In contrast to fWHR, these two facial ratios incorporate the length and breadth of the jaw—an area of the face with a long history of research in biological anthropology (Lundström & Lysell, 1953; Merton & Ashley-Montagu, 1940;), clear sexual dimorphism across populations (Franklin et al, 2008; Saini et al., 2011), associations with other SSCs (Hodges-Simeon et al., 2016), and known associations with age and testosterone during development (Hodges-Simeon et al., 2016; Snodell et al., 1993; Verdonck et al., 1999). Further, we include three variants of fWHR used in the literature: fWHR*nasion*, fWHR*brow*, and fWHR*stomion* (see Fig 1 and Table 1 for a guide to the facial ratios used in the present research and in previous studies). We use this specific terminology here to increase clarity, as each of these variants has separately been termed “fWHR” in the literature. Researchers have largely treated these variants as interchangeable, yet it is unclear whether this decision is justified—i.e., to what extent the variants overlap with one another.

Finally, in all analyses, we control for individual differences in facial adiposity using BMI (Coetzee et al., 2010). Lefevre et al. (2013) found sexual dimorphism in fWHR disappeared after controlling for BMI. A meta-analysis of studies before 2015 indicated that higher BMI was associated with larger fWHRs in adults (Geniole et al., 2015), yet only a third of the studies reviewed for this paper control for individual differences in adiposity (see Table 1). This may also be an important control in behavioral research; for example, Deanor et al. (2012) identified body weight (which likely overlaps muscle mass), not fWHR, as a predictor of aggression among athletes (see also Mayew, 2013).

**Fig 1.**
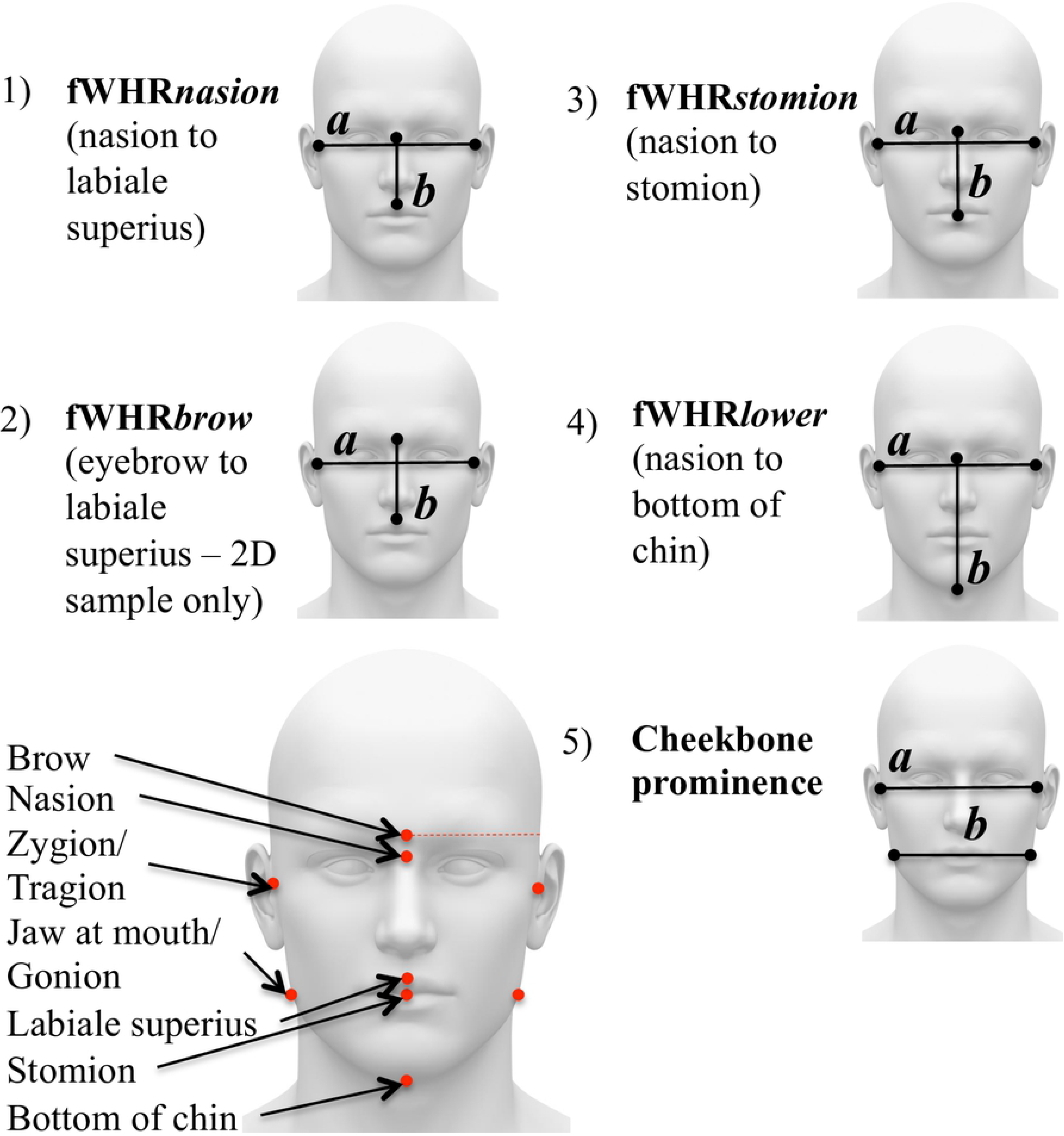
Candidate facial masculinity ratios used in the present research

**Table 1.**
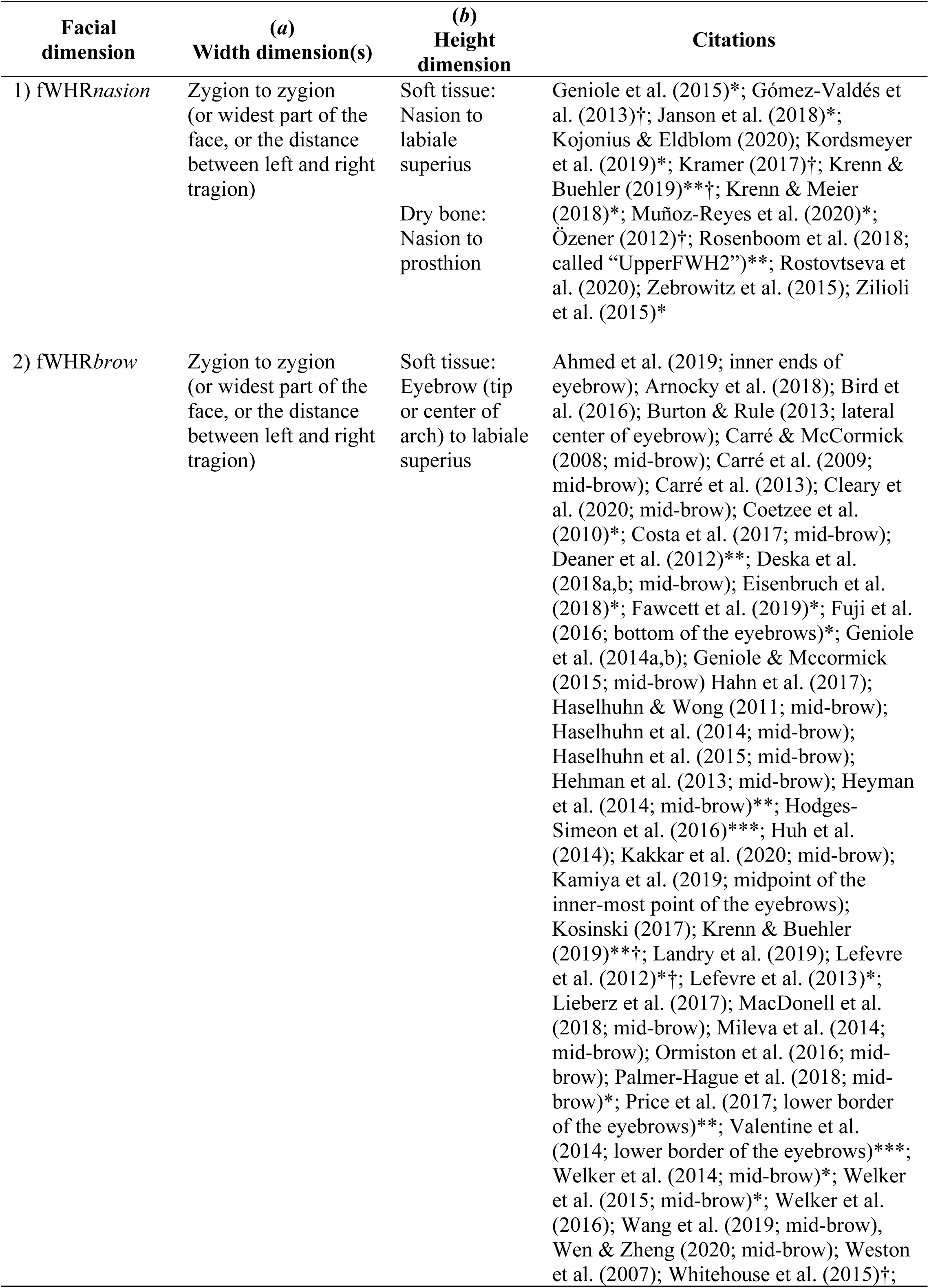
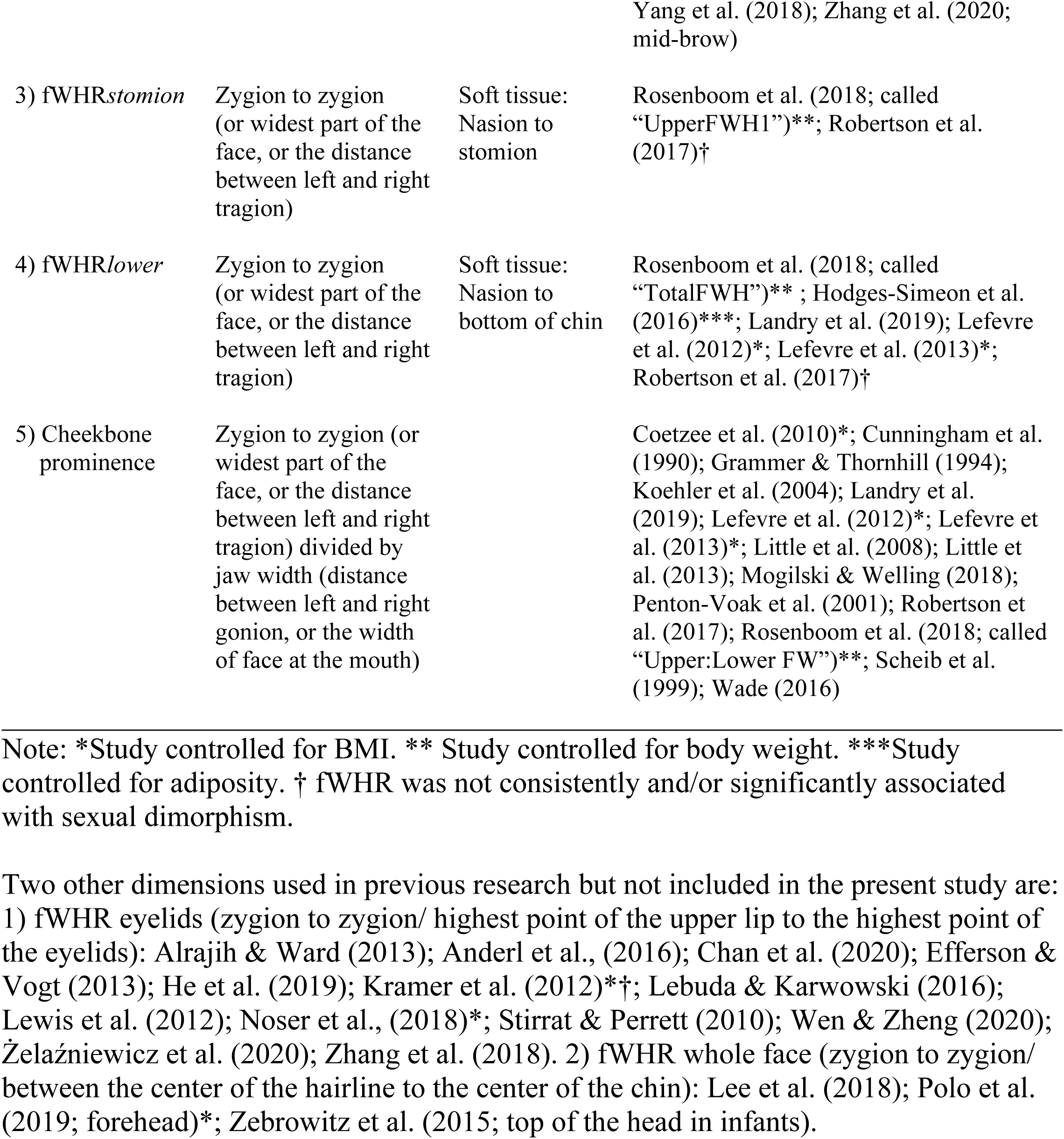
Facial ratios used in the present research

## Methods

### 3D European/Caucasian Sample

#### Participants

3D facial scans were obtained from the 3D Facial Norms data set (see Weinberg et al., 2017 for a detailed sample description). Participants were recruited from four US cities (Pittsburgh, Seattle, Houston, and Iowa City), primarily through target advertisements. Only individuals who had no history of craniofacial trauma, congenital malformations, or facial surgery were permitted to participate (Kesterke, et al., 2016).

The sample consisted of 2,449 unrelated individuals of European-Caucasian ancestry between the ages of 3-40 (1502 females and 952 males). Individuals were classified into four age groups: child (3-6 years of age, *N* = 193), juvenile (7-11 years of age, *N* = 199), adolescent-to-young adult (12-21 years of age, *N* = 580), adult (22-40 years of age, *N* = 1477). We classified ages 19-21 as “adolescents” for several important reasons. First, the end of adolescence is ambiguous and variable across individuals and populations. Western societies arbitrarily set this at 18; however, life history theory marks the end of adolescence with the end of growth and birth of first offspring—events that may vary widely. Second, while male adult height may be reached in the late teens (but not always; Bogin, 1999), growth in other tissues (i.e. muscle mass) often continues after age 18 (Schutz et al., 2002). Third, endocrine maturation (i.e. rapidly increasing production of sex steroids) usually continues into the early 20s for males (Butler et al., 1989; Elmlinger et al., 2004; Kelsey et al., 2014; Hodges-Simeon et al., 2015). Therefore, development of T-mediated traits will also likely extend past age 18.

#### Instruments

Digital stereophotogrammetry was used to obtain 24 landmark distances from the 3D facial scans, from which 5 were used in the present study (nasion, labiale superius, stomion, bottom of the chin, and tragion as a proxy of zygion; see Fig 1). We also utilized two additional distances collected with direct anthropometry using spreading calipers (GPM Switzerland): maximum facial width (zygion to zygion) and mandibular width (gonion to gonion). Previous investigations have verified that data collected from facial images using digital stereophotogrammetry are highly replicable and precise (Aldridge, Boyadjiev, Capone, DeLeon, & Richtsmeier, 2005); nevertheless, we examined correlations between fWHR measures calculated using facial width from landmark distances versus direct anthropometry. All were highly correlated: fWHR*nasion* (*r* = .92), fWHR*stomion* (*r* = .91), fWHR*lower* (*r* = .89), and cheekbone prominence (*r* = .87). All models described in the results were also run using the caliper-derived ratios, which altered Beta values by only trivial amounts.

#### Facial landmarks and masculinity ratios

Facial width was measured from the left to the right tragion, the point marking the notch at the superior margin of the tragus, where the ear cartilage meets the skin of the face. The upper boundary of facial height was measured from the approximate location of the nasion, the midline point where the frontal and nasal bones contact. The lower boundaries for mid-facial height included the labiale superius, the midline point of the vermilion border of the upper lip at the base of the philtrum (for fWHR*nasion*); the stomion, the midpoint of the labial fissure (fWHR*stomion*); and the bottom of the chin (fWHR*lower*). See Fig 1 and Table 1. Ratios were computed by dividing facial width by facial height; greater fWHRs reflect relatively wider faces relative to the height dimensions. Cheekbone prominence was a ratio of facial width to mandibular width. In this sample, mandibular width was measured using a caliper at the left and right gonion. Previous research on cheekbone prominence in front-facing 2D photographs has approximated this location (Lefevre et al., 2012) or used the width of the face at the mouth (Hodges-Simeon et al., 2016; Penton Voak et al., 2001). Information about the location of the brow was not available in the 3D renderings; therefore, of the ratios shown in Fig 1, fWHR*brow* could not be used with the 3D sample.

Ratios (rather than measures of individual facial dimensions) are often utilized in previous research for several reasons. First, for 2D photographs in particular, ratios offer greater ease of measurement; that is, no corrections are necessary for distance from the camera, ontogenetic scaling, or deviations from the Frankfurt plane. Second, because of this ease, ratios have been increasingly adopted in disciplines outside of biological anthropology; as such, there is now a growing literature of fWHR results that require evolutionary and ontogenetic explanation.

#### Anthropometrics

Self-reported height and weight were collected from each participant, and then used to calculate BMI. See www.facebase.org/facial_norms/notes/ for more information on the sample.

### 2D Bolivian Tsimane Sample

#### Population

The Tsimane are a small-scale, kin-based, group of hunter-horticulturalists who reside in the Amazonian lowlands of Bolivia. They obtain relatively few calories from market sources, have little access to modern medicine, and experience high rates of infectious diseases (Gurven, Kaplan, Winking, Finch, & Crimmins, 2008; Martin et al., 2012; Vasunilashorn, Crimmins, Kim, Winking, Gurven, Kaplan, & Finch, 2010). On average, individuals experience high rates of infection; for example, approximately 60% of individuals carry at least one parasite (Vasunilashorn et al., 2010). As such the Tsimane experience high rates of chronic inflammation, characteristic of populations living in environments with high pathogen loads (Gurven et al., 2008).

#### Participants

Participants consisted of 139 peripubertal individuals (73 males and 66 females) between the ages of 7 and 21. Participants’ ages were estimated by comparing their self-reported age to their age taken from the Tsimane Health and Life History Project (THLHP) census (Gurven, Kaplan, & Supa, 2007). When there was a discrepancy between participants’ self-reported and census ages, census age was used (see Hodges-Simeon et al., 2013, for further explanation of age estimation methods). Following our 3D sample, participants were divided into juvenile (age 7 to 11) and adolescent (age 12 to 21) age groups.

#### Facial measurement

To obtain facial measurements, we first took high-resolution, front-facing color photographs of participants using a 12MP Sony camera. Participants’ heads were positioned along the medial-sagittal plane and they were instructed to have a neutral facial expression. Eleven trained research assistants (RAs), from Boston University and University of California Santa Barbara, placed landmarks on all facial photographs using the image-editing software GIMP and each photograph was processed by three RAs. The research assistants were blind to the hypotheses of the researcher and did not know any of the photographed individuals. The research assistants recorded the x-y coordinates for each landmark of the face twice. The coordinates were averaged (i.e., a total of six x coordinates and six y coordinates per landmark) to establish final landmark coordinates (α = .88, for males, α = .98 for females for the entire sample). Feature measurements were standardized using inter-pupillary distance. Landmarks of interest and ratios are shown in Fig 1. fWHR*nasion,* fWHR*stomion*, and fWHR*lower* were calculated based on the same landmarks as described for the 3D sample above. Because the location of the nasion must be approximated in soft tissue (the nasion is the midline point where the frontal and nasal bones contact), we anticipate more error for this point. fWHR*brow* was calculated in the same way as in Carré & McCormick (2008): bi-zyomatic breath was divided by height of the face from the top of the lip to the middle of the brow. Cheekbone prominence was a ratio of facial width to the width of the face at the mouth (Hodges-Simeon et al., 2016; Penton Voak et al., 2001).

#### Anthropometrics

Standard anthropometric protocols were used to assess growth and energetic status (Lohman et al. 1988); participants wore light clothing and no shoes for measurement of height and weight (to determine BMI).

#### Data Screening and Analysis

SPSS 24 was used for all analyses. To correct for small deviations from normality all study variables were log-transformed. Although transformation only altered results by trivial amounts, we report results here using the transformed variables. All assumptions for multivariate analysis (i.e., multi-collinearity, normality, linearity, and homogeneity of variance) were met. Variance Inflation Factors (VIFs) were used to assess multicollinearity; all VIFs < 2.

For analyses, alpha level was set at 0.05 (two-tailed). As a first step, we examined bivariate correlations between all pairs of variables. Point biserial correlations were examined for associations between sex and all other variables of interest (see Table 2). We employed correlations to assess the degree of multicollinearity among different measures of fWHR. Inspection of correlations between different measures of fWHR revealed only small differences across the age groups (i.e., fWHRnasion and fWHRstomion were closely correlated regardless of the age category). Therefore, in the interest of reducing the number of tests, we collapsed across age categories to examine correlations for males and females separately, controlling for age (see Supplement for Table S1 for the 3D sample and Table S2 for the 2D sample). We then proceeded to conduct standard (i.e., simultaneous) multiple regressions, within each face set and age group (Table 3).

In both samples, males were coded “1” and females were coded “2”; therefore, in the results presented below, positive associations with sex indicate that female means are higher on this trait. Given the importance of accurate coding of sex for the interpretation of results, we examined the association between sex and height—a known SSC—in both samples. In the 3D sample, sex was inversely correlated with body height in adults (*r* = -.71, *p* < .001) and in adolescents (*r* = -.50, *p* < .001), with adult males showing the expected height advantage over females. Among adolescents in the 2D sample, sex was inversely correlated with height but did not reach conventional levels of significance (*r* =-.26, *p* = .08); therefore, we examined the association between sex and voice pitch (data from Hodges-Simeon et al., 2013), which is more strongly dimorphic than height (Puts et al., 2012). Sex was positively correlated with voice pitch controlling for age (*r* = .46, *p* < .001). That is, being female was associated with higher voice pitch, which confirms accurate sex coding in the 2D sample.

Curve Expert Version 1.5.0 was used to determine a best-fit algorithm for patterns of age-related change in facial masculinity ratios. Goodness-of-fit was assessed using the coefficient of determination (*R*^2^). In Hodges-Simeon et al. (2013, 2016), these methods were used to demonstrate evidence for growth spurts in height and voice pitch.

## Results

### Correlations

**Table 2.** Point-biserial correlations with sex across age groups (positive values indicate that females are larger)

|  | **fWHR-**<br>**nasion**<br>(nasion<br>to labiale<br>superius) | **fWHR-**<br>**brow**<br>(brow to<br>labiale<br>superius) | **fWHR-**<br>**stomion**<br>(nasion<br>to<br>stomion) | **fWHR-**<br>**lower**<br>(nasion<br>to<br>bottom<br>of chin) | **Cheek-**<br>**bone**<br>**Promin-**<br>**ence** | **BMI** | **Height** | **Age** |
| --- | --- | --- | --- | --- | --- | --- | --- | --- |
| **<u>3D Sample</u>** |  |  |  |  |  |  |  |  |
| **Adults** | -.04 | n/a | -.08*** | .07* | .08** | -.10*** | -.71*** | .07† |
| **Adolescents** | -.01 | n/a | -.04 | .02 | .04 | .02 | -.50*** | .15*** |
| **Juveniles** | -.10 | n/a | -.12 | -.12 | -.12 | -.08 | .01 | .09 |
| **Children** | -.10 | n/a | -.13 | -.12 | -.05 | -.02 | -.01 | .03 |
| **<u>2D Sample</u>** |  |  |  |  |  |  |  |  |
| **Adolescents** | .18† | -.44** | -.04 | .10 | -.11 | .15 | -.26† | .14 |
| **Juveniles** | .24 | -.43*** | .09 | .32† | -.09 | -.03 | .01 | .33† |

Significance levels (two-tailed): † $P < 0.10$ , \* $P < 0.05$ , \*\* $P < 0.01$ , \*\*\* $P \leq 0.001$ .

*3D European/Caucasian sample.* Point-biserial correlations revealed significant sex differences (positive values indicate females are larger) in fWHR*stomion* (*r* = -.08, p = .001), fWHR*lower* (*r* = .07, p = .001), cheekbone prominence (*r* = .08, p = .001), and BMI (*r* = -.10, p = .001) in adults, but not adolescents, juveniles, and children (see Table 2). Age was correlated with sex in both adults (*r* = .07, p = .01) and adolescents (*r* = .15, p < .001), underscoring the need to control for age in further analyses. Collapsing across age groups (and controlling for age), fWHR*nasion,* fWHR*stomion*, and fWHR*lower* showed high collinearity given their shared points of measurement (*r*s = .78-.96; see Table S1 for exact values). For males and females, cheekbone prominence was moderately associated with fWHR*nasion* (*r* = .15 and .07, respectively), fWHR*stomion* (*r* = .17 and .09), and fWHR*lower* (*r* = .18 and .12). BMI was positively associated with fWHR*nasion,* fWHR*stomion*, and fWHR*lower* for both males and females, indicating increased facial width with increasing BMI. Cheekbone prominence was inversely associated with BMI for females only, indicating that weight gain affects the breadth of the lower face for females. See Table S1.

*2D Bolivian Tsimane sample.* Males had larger fWHR*brow* (*r* = -.44, p = .001, in adolescents; *r* = -.43, p = .001, in juveniles) and fWHR*lower* (r = -.29, p = .004, in adolescents), but there were no sex differences in fWHR*nasion* and cheekbone prominence. See Table 2. We also looked at the relationships between fWHR measures to explore the extent to which these measures co-varied. fWHR*nasion* and fWHR*stomion* were correlated in males (*r* = .71, *p*<.001) and females (*r* = .40, *p* < .001), similar to the 3D sample. fWHR*brow* was also closely associated with fWHR*nasion* (*r* = .82, *p* < .001 and *r* = .78, *p* < .001) and fWHR*stomion* for males and females, respectively. Cheekbone prominence was significantly associated with fWHR*lower* (*r* = .63, *p* < .001 and *r* = .31, *p* < .01). In contrast to the 3D sample, fWHR*lower* was not significantly associated with fWHR*nasion*; however, fWHR*lower* was correlated with fWHR*brow* (*r* = .40, p < .001 and *r* = .61, *p* < .001) and fWHR*stomion* (*r* = .50, *p* < .001 and *r* = .79, *p* < .001). Also in contrast to the 3D sample, cheekbone prominence was inversely correlated with fWHR*nasion* (*r* = -.40, *p* < .001 and *r* = -.39, *p* < .01) and uncorrelated with fWHR*brow* and fWHR*stomion*. See Table S2.

### Are fWHR and/or other commonly used masculinity ratios sexually dimorphic in adults?

*3D European/Caucasian sample.* Zero-order correlations indicated that both BMI and age were associated with sex; therefore, we employed multiple regression to examine the effects of sex on facial masculinity ratios while controlling for these potential confounds. Four separate multiple regression models were employed with sex, age, and BMI as predictors and fWHR*nasion*, fWHR*stomion*, fWHR*lower*, and cheekbone prominence as the outcome variables (see Table 3). Sex was a significant predictor of fWHR*stomion* (*ß* = -.05, *p* < .05), fWHR*lower* (*ß* = .09, *p* < .001), and cheekbone prominence (*ß* = .08, *p* < .01), but not fWHR*nasion (ß* = -.01, *p* = .84). In other words, males showed the expected pattern of larger mandible breadth (i.e., smaller cheekbone prominence) and longer chin (i.e., smaller fWHR*lower*). Males showed significantly wider faces relative to the midface, but only when the midface extended to the stomion (i.e., fWHR*stomion*), and not when it terminated at the labiale superius (fWHR*nasion*). This finding was surprising given the shared variance in fWHR*nasion* and fWHR*stomion* (*r* = .96; see Table S1). Post-hoc analyses showed a significant sex difference in upper lip height in this sample (*ß* = -.38, *p* < .001) controlling for age and BMI; that is, males have significantly larger upper lip height than females.

BMI was a significant predictor of the outcome variables in all models. Age was also a significant negative predictor for fWHR*nasion* and fWHR*stomion*; as individuals age from 22 to 40 years, both of these fWHR measures get smaller, likely reflecting a lengthening of the midface with aging (see Table 3). See also Fig 2 for visual representation of changes in the variables of interest with age.

**Table 3.**
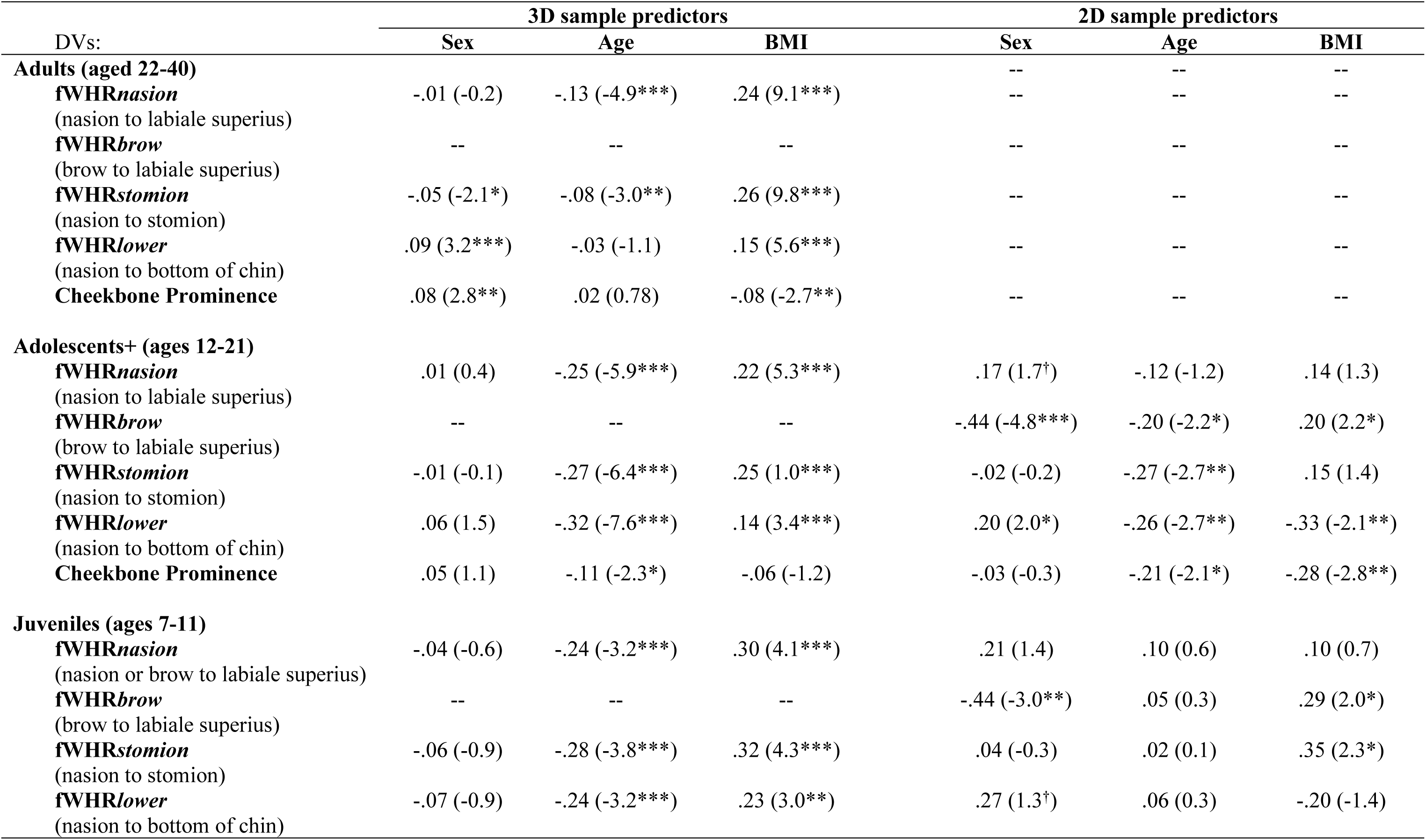
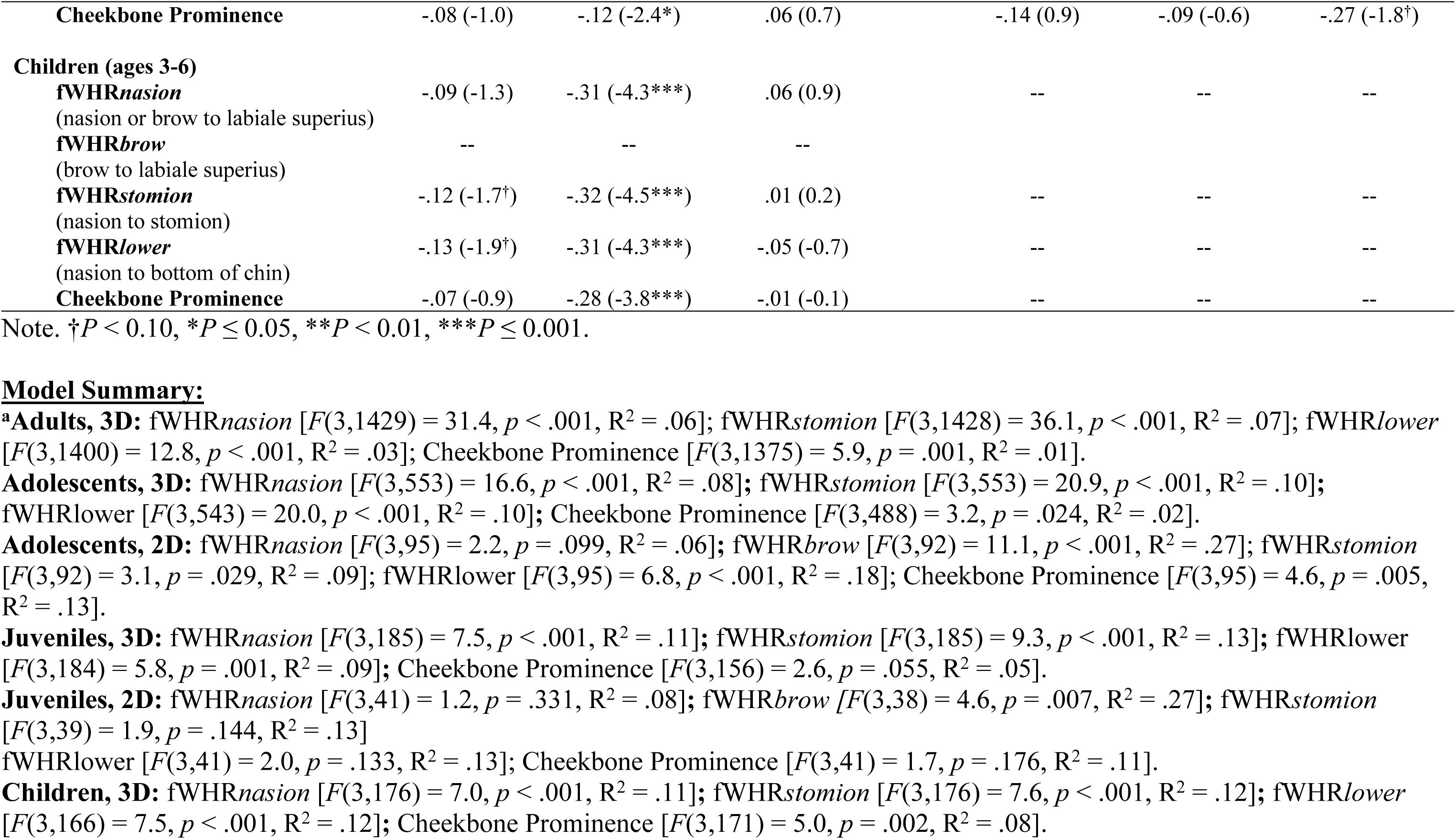
Multiple regression models. Standardized Beta coefficients shown with *t* statistic in parentheses. Positive values for sex indicate female ratios are larger.

**Fig 2.**
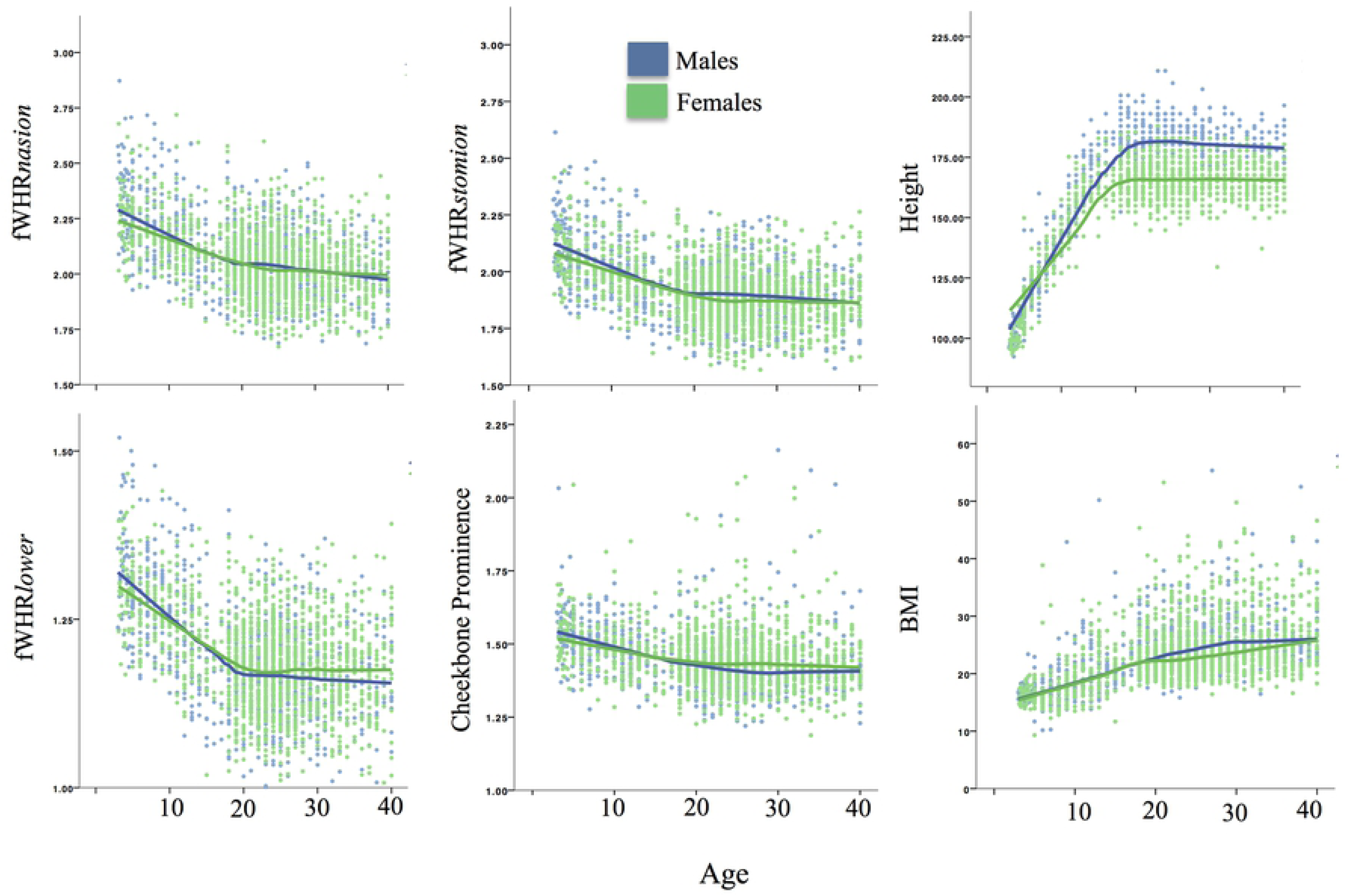
Facial masculinity ratios, height, and BMI by age and sex (3D sample)

### Are fWHR and/or other commonly used masculinity ratios sexually dimorphic in sub-adults?

*3D European/Caucasian sample.* Separate multiple regression models were again conducted for each age group—children, juveniles, and adolescents— and paralleled those for adults. Across all sub-adult age groups, sex was not a significant predictor of any of the masculinity ratios while age was a significant inverse predictor of all facial ratios (see Table 3 for standardized Betas and *t* statistics). With sub-adult growth, fWHR*nasion* (*ß* = -.25, *p* < .001) and fWHR*stomion* (*ß* = -.27, *p* < .001) became smaller—facial width decreased relative to midface height (i.e., became less masculine based on current conceptualizations of fWHR). fWHR*lower* (*ß* = -.32, *p* < .001) and cheekbone prominence (*ß* = -.11, *p* < .05) also became smaller, indicating childhood growth in mandible dimensions relative to bizygomatic width. Similar to the adults, BMI was a significant positive predictor of fWHR*nasion*, fWHR*stomion*, and fWHR*lower* in juvenility and adolescence but not childhood (*ß*s = .14 - .32; see Table 3). In other words, juveniles/adolescents with greater somatic adiposity (and, by extension, facial adiposity) had wider faces relative to facial height. See Table 3.

*2D Bolivian Tsimane sample.* Because brow information was available for the 2D sample but not the 3D sample (see Methods for more information), we examined multiple regression models predicting fWHR*brow* as well as the other 4 ratios. In adolescents, sex was a significant negative predictor of fWHR*brow* (*ß* = -.44, *p* < .001), but not fWHR*stomion* or fWHR*nasion*, for which sex approached significance as a *positive* predictor (*ß* =.17, *p* = .09). Again, these results were surprising because fWHR*brow* and fWHR*nasion* were correlated with each other (*r* = .82, *p* < .001). Post-hoc analyses were employed to determine if the distance from the nasion to the brow was sexually dimorphic and could be driving the opposing relationships with sex. Controlling for age and BMI, sex was a very strong predictor of nasion-to-brow distance (*ß* = .72, *p* < .001), with females having higher-placed brows relative to the nasion position. A similar pattern was found for juveniles (*ß* = .75, *p* < .001; see Table 3), indicating this sex difference is present prior to puberty. See Fig 3 for nasion-to-brow distance by age.

**Fig 3.**
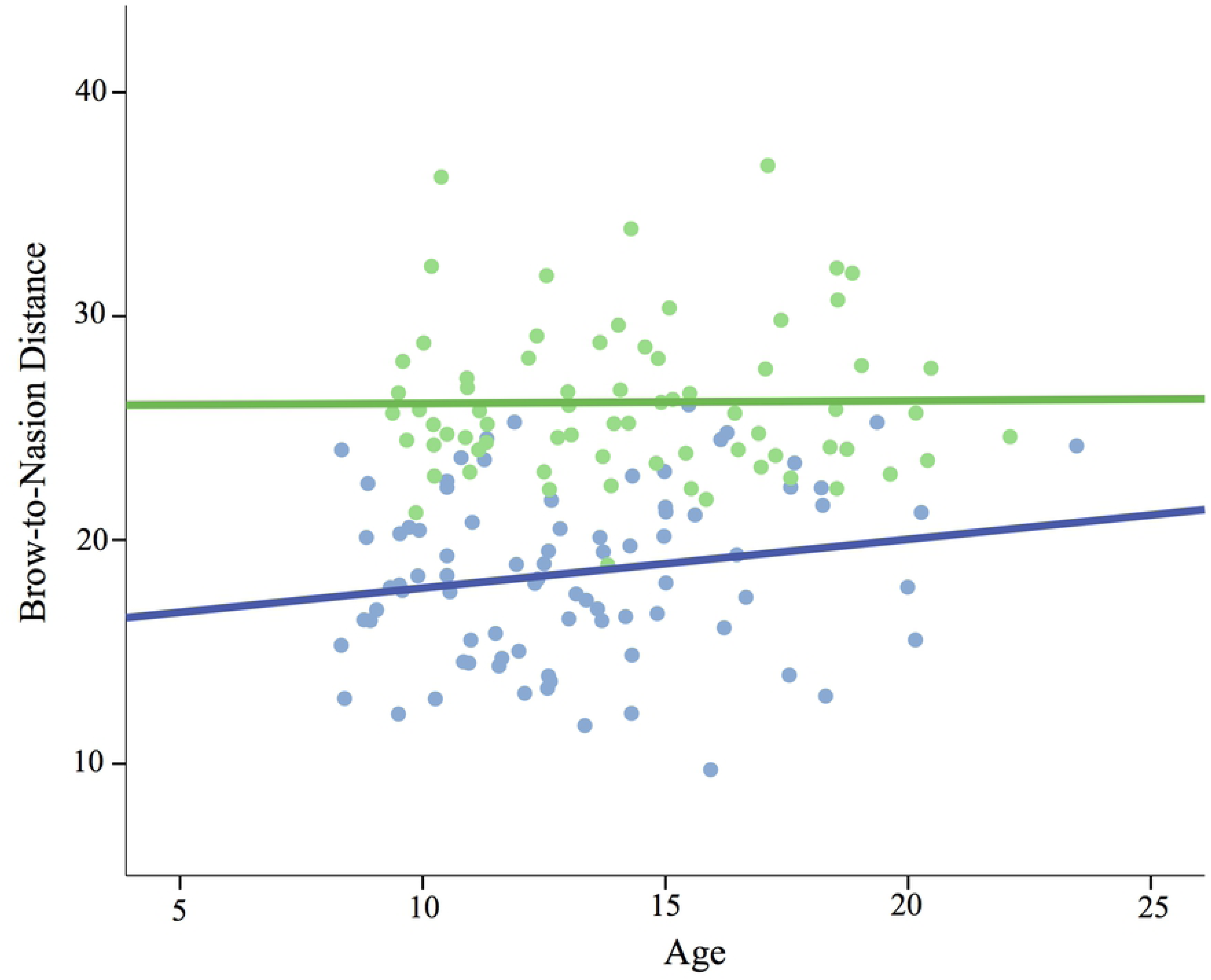
Brow-to-nasion distance by age and sex (2D sample)

Results also showed that sex was a significant positive predictor of fWHR*lower* in adolescents (ß = .20, *p* = .04) and approached conventional significance in juveniles (ß = .27, *p* = .08).

### What is the pattern of sex-specific ontogeny for facial masculinity ratios?

*3D European/Caucasian sample.* Because analyses thus far showed a significant effect of age on facial ratios across age groups, we explore age-related changes by sex in Fig 2. Visual inspection of results indicates declining facial width relative to height during sub-adult growth as well as during adulthood, supporting conclusions about the effects of age drawn from regressions above.

In order to assess the extent to which facial masculinity ratios exhibit changes in velocity during adolescence—i.e., a growth spurt—we examined whether a sigmoidal model explained more variance than a linear one. Because fWHR*stomion*, fWHR*lower*, and cheekbone prominence were found to be sexually dimorphic in adulthood, the pattern of development for each of these ratios was examined for evidence of a growth spurt. As in Hodges-Simeon et al. (2016), we found no evidence of changes in facial ratio growth velocity during adolescence.

Visual inspection of the scatterplots suggested that fWHR*lower* might become sexually dimorphic in later adolescence; therefore, post-hoc analyses were also conducted to determine if restricting the age range to over 14 in both samples changed the results for the adolescent age group. In the 3D sample, fWHR*lower* was sexually dimorphic (*ß* = .11, *p* =.02) among those aged 14 to 21. Restricting the age range did not change the effect of sex for any of the other ratios. In the 2D sample, restricting the age range to 14+ did not substantially change the results; however, fWHR*nasion* did reach conventional levels of significance (*ß* = .16, *p* =.049). That is, over-14 female adolescents had significantly larger fWHR*nasions* than did males.

## Discussion

The goal of the present research was to address ongoing debates on the existence and evolutionary origins of sex-typical variation in fWHR and other facial masculinity ratios using ontogenetic evidence. We examined sex differences in five different ratios across sub-adult and adult age groups in 2D photos and 3D renderings in two distinct populations. Results showed that 3 variables predict significant variation in facial masculinity ratios—sex, age, and BMI. Each reveals potentially important clues to inconsistencies in past fWHR research and suggest agendas for future research.

### Summary of results

First, sex was a significant predictor of some but not all facial masculinity ratios. Across both samples, those ratios that incorporated dimensions of the lower face—i.e., the length (fWHR*lower*) and breadth (cheekbone prominence) of the mandible—suggest a history of sexual selection. In the adult 3D sample (ages 22 to 40), fWHR*lower* and cheekbone prominence were clearly sexually dimorphic, with males again showing a longer (in terms of fWHR*<u>lower</u>* where jaw size augments length) and wider (in terms of cheekbone prominence where jaw size augments width) lower face than females. fWHR*lower* also showed the expected ontogenetic pattern for SSCs; that is, sexual dimorphism developed in the life stage following puberty. In the 2D sample, among adolescents (aged 12 to 21), but not among juveniles (aged 7 to 11), sex was a significant predictor of fWHR*lower*. In the 3D adolescent sample (aged 12 to 21), sex differences were not found; however, when the age group was restricted to later adolescent ages— i.e., 14 to 21—a significant sex difference emerged, suggesting that lower face development may occur later in adolescence. These findings accord with a long history of research in biological anthropology showing differential growth in the mandible among male *Homo sapiens* (Enlow & Harris, 1964; Lundström & Lysell, 1953; Merton & Ashley-Montagu, 1940), which produces measureable sex differences across diverse populations (Claes et al., 2012; Matthews et al., 2018). These findings also make sense in light of research showing associations between fWHR*lower* and baseline testosterone levels (Hodges-Simeon et al., 2016), one testosterone-related genetic variant (Roosenboom et al., 2018), as well as other testosterone-dependent traits, like upper body strength (Hodges-Simeon et al., 2016).

Our review of the literature, although not exhaustive, showed substantial variation in the way fWHR is measured when the midface is used as the height dimension (see Table 1). Facial width is relatively consistent across studies; however, midface height has several variants, which we called fWHR*nasion*, fWHR*brow*, and fWHR*stomion* (see Fig 1). Despite high correlations among these measures, sex differences in these variants were not consistent across measures and samples. In the 3D sample, fWHR*stomion* was larger in adult males, yet closely correlated fWHR*nasion* was not dimorphic. Post-hoc analyses showed that this pattern of results was driven by greater upper lip height in males compared with females (also found by Kesterke et al., 2016; Matthews et al., 2018). Sexual dimorphism in upper lip height illustrates that variants of fWHR should not be treated as interchangeable in research. In the 2D sample, fWHR*stomion* was not dimorphic, while fWHR*nasion* was significantly larger in *females* rather than males (among those over 14). It is possible that variation across these samples may be due to inter-population differences in the presence and degree of sexual dimorphism in fWHR; for example, Kramer et al. (2017) found significant sex differences in fWHR*nasion* among East Asian populations but not any other groups. The degree of SSC development may vary with energetic stress (Hodges-Simeon et al., 2013) and greater sexual dimorphism has been found among energy-abundant societies (Stinson, 1985), underscoring the need to sample across a range of diverse human socioecologies, as we have done here.

Our 2D sample included landmarks on the eyebrow, which was not available for the 3D renderings. fWHR*brow* was sexually dimorphic, with males showing the expected wider faces relative to females. Again, this was surprising because closely correlated fWHR*nasion* and fWHR*stomion* were not dimorphic. Post-hoc analyses revealed that the distance from the nasion to the brow accounts for this pattern of results, with females showing substantially higher brows than males. Like mandible size, this finding accords with previous research on greater supraorbital, or brow ridge, size in male *Homo sapiens* (Claes et al., 2012; Gavin & Ruff, 2012; Shearer et al., 2012), which is likely associated with lower-set eyebrows. Work in growth modeling has shown that males’ brow ridge grows faster during adolescence, giving rise to observable sex differences by age 16 (Matthews et al., 2018).

Our results also showed that sexual dimorphism in fWHR*brow* emerges early, with sex being a significant predictor even in our juvenile sample. The ontogeny of secondary sexual traits is traditionally characterized by differential male and female growth arising from sex steroid hormone increases in puberty (Ellison, 2012; Hochberg, 2012). These findings, however, suggest that certain sexually dimorphic face features may diverge prior to puberty—in other periods characterized by hormonal switch points (i.e., prenatal, early post-natal, post-adrenarche). This conclusion is supported by a number of studies that have identified significant early-life sex differences in the face (Bulygina et al., 2006; Weinberg et al., 2014; Whitehouse et al., 2016) and other aspects of the phenotype (e.g., Fouquet et al., 2016). Matthews et al. (2018) observed that there were two phases in the emergence of facial sexual dimorphism—ages 5 to 10 (i.e., the post-adrenarche period; Campbell, 2011) and ages 12 onwards. Some aspects of facial sexual dimorphism were present in the first phase and became more exaggerated in the second phase (i.e., forehead, chin, and cheeks), whereas others did not emerge until the second phase (i.e., nose, brow ridge, and upper lip). Sexual dimorphism in several other SSCs begins before puberty; for example, human female infants show greater body fat from birth onwards (Koo et al., 2000). The ultimate reasons for different emergence patterns should be addressed in future research; however, one interpretation is that mating and status competition may begin before puberty in humans.

A lower brow position may be an important factor in raters’ perceptions of aggressiveness, fighting ability, masculinity, dominance, and threat in those with high fWHR*brow* (Geniole et al., 2015; Geniole & Mccormick 2015; Zilioli et al., 2015). Research on emotion attribution from facial features has shown that lower-placed eyebrows are perceived as more threatening and aggressive regardless of the facial expression and that raters have greater anger recognition accuracy for high fWHR faces and greater fear accuracy for low fWHR faces (Deska et al., 2017, which used brow position). Further, faces where the chin is tilted forward or backward have higher fWHR and are perceived as more intimidating as a result (Hehmen et al., 2013, which also used the brow). Lower brow position in males may be a cause or consequence of the evolution of the anger expression and head orientation; that is, sexually dimorphic attributes may have co-evolved with universal facial expressions of anger and fear (Sell et al., 2014).

### Confounds in fWHR research: Age and BMI

Across both samples, age was a significant inverse predictor of fWHR measures, controlling for sex and BMI. In the 3D sample, age was a consistent negative predictor of facial masculinity ratios from age 3 to adulthood; however, the effect was more pronounced in sub-adult groups. In other words, the face becomes less wide relative to midface height, lower face height, and chin breadth throughout childhood growth, i.e., less “babyfaced” (Zebrowitz et al., 2015). This is likely a consequence of the decreasing relative size of the cranial vault from birth to adulthood along with increases in nose and mandible growth (Matthews et al., 2018). In addition, the 3D sample showed that fWHR*nasion* and fWHR*stomion* continue to decrease with adulthood ageing, which has been shown in previous research (Hehman et al., 2014; Robertson et al., 2017), although the slope is not as steep as among sub-adult groups (see Fig 2). This effect may be due to age-related collagen degradation (Yasui et al., 2013) and/or changes in the bony structure (Shaw et al., 2011). Overall, these findings point to age as an important variable to consider in sample selection and data analysis in fWHR research.

BMI was also a significant predictor of most fWHR measures across juvenile, adolescent, and adult age groups (see Table 3). BMI was used as a proxy measure for fat stores and controlled in all analyses because fat tends to be deposited on the cheeks and chin, increasing facial width. Previous research has consistently shown that BMI is correlated with a higher fWHR (Geniole et al., 2015); yet a minority of studies reviewed for this paper control for it (see Table 1). The role of BMI in predicting individual differences in facial masculinity ratios speaks to the importance of examining fWHR in both dry bone and soft tissue faces. Evidence suggests that there may be differential selection on bone and fat/muscle in humans and that each may separately contribute to increases in fWHR. For example, in one forensic sample, men with lower fWHRs were significantly more likely to die from contact violence than were men with higher fWHR, suggesting that men with relatively wider faces were more likely to survive aggressive encounters with other men (Stirrat et al., 2012). The authors hypothesized that greater zygomatic buttressing may have benefited ancestral men by reducing the negative effects of craniofacial impact. Yet measures of fWHR from 2D photographs cannot distinguish facial breadth due to bony dimensions, which are more substantial in men, versus fat deposits, which tend to be greater in women (Lassek & Gaulin, 2009). Previous studies have shown that the cheek region is sexually dimorphic (Matthews et al., 2018) and our results showed that BMI affects cheekbone prominence in females but not males. Finally, little research has considered how sex differences in facial muscle may impact fWHR dimensions; one recent study showed that the brachyfacial face type, which overlaps with high fWHR, has greater masseter volume than more narrow face types (Woods & Wong, 2016).

### Ontogeny and sexual selection

The broader goal of this research was to emphasize the importance of using ontogenetic data to address questions in sexual selection research, using fWHR as a model case. We point to four questions that may be asked of this type of data that should corroborate conclusions drawn from data on adults, providing a roadmap for future researchers to use developmental patterns to substantiate claims about sexual selection pressures. First, do sex differences arise in coordination with the onset of mate competition? Second, do sex differences arise from differential male or female growth? Third, does the purported sexually selected trait exhibit a spurt? And finally, do these traits co-vary with sex steroid hormones and/or other SSCs? Our results show that only fWHR*lower* exhibits the expected pattern of ontogeny for a sexually selected male trait

As a further example of a SSC with a clearer history of sexual selection, we point to research on the low human male voice. During puberty, increased production of testosterone causes males’ vocal folds to thicken and their larynxes to descend, producing a lower pitched and more resonant sounding voice (Abitbol, Abitbol, & Abitbol, 1999; Butler et al. 1989; Fitch & Giedd, 1999; Harries et al., 1998). Male adolescents experience a decrease in fundamental and formant frequencies, which jointly contribute to perceived lower pitch, as their vocal folds thicken and lengthen. This decrease happens in a “spurt” (Hodges-Simeon et al., 2013). By adulthood, the sex difference in fundamental frequency is over 5 standard deviations (Puts et al., 2011). Lower pitched voices are rated as more attractive-sounding by women and more dominant-sounding by both sexes (Feinberg et al., 2005; Jones et al., 2010; Puts et al., 2007). Furthermore, in one natural fertility population, men with lower pitched voices were found to father more offspring (Apicella, et al., 2007). Finally, sexually dimorphic vocal parameters are correlated with body size (Pisanki et al., 2014), muscle mass during adolescence (Hodges-Simeon et al., 2014), and aggressiveness (Puts et al., 2011). These various sources of evidence jointly lend greater confidence to the assertion that male vocal traits are SSCs.

## Limitations

This research has several limitations. We sought to compare the pattern of fWHR ontogeny in two distinct populations (European-decent Caucasians and indigenous-decent Bolivians); however, there were methodological differences between the two that prohibit a direct comparison. First, besides being 3D and 2D respectively, landmarks were placed by a different set of researchers, which could have introduced bias. Further, cheekbone prominence was measured using a caliper distance in the 3D sample and a landmark distance in the 2D sample, based on what was available in the datasets. Further research is needed which directly compares across populations using the same methodology (see Kramer, 2017). Second, the nasion landmark was used in Weston et al. (2007)’s original research on facial width in dry bone samples; however, it should be used with caution in soft tissue studies. The nasion refers to the midline point where the frontal and nasal bones contact (i.e., the nasofrontal suture). Although informed by previous research (Kolar & Salter, 1997), this exact position poses more of a challenge in soft tissue photos or renderings; therefore, there may be a larger degree of error in this landmark. Our results suggest that when fWHR is measured in soft tissue, brow position should be used rather than the nasion. Finally, this research highlights the importance of age, yet the data are cross-sectional. Future studies on intra-individual longitudinal change would help clarify the effect of age and BMI on sex differences in fWHR.

## Conclusions

These findings add an ontogenetic perspective to the ongoing debate on the history of sexual selection on fWHR. Our results show that only fWHR*lower* exhibits the classic pattern of ontogeny for a sexually selected human male trait —i.e., adult sex differences in fWHR*lower* along with greater lower-face growth in males relative to females during adolescence. These findings also highlight potential confounds that may be responsible for inconsistent findings in the fWHR literature (i.e., age—due to both sub-adult growth and adult ageing—and BMI), and also reveal via post-hoc analysis some features (brow position and lip height) that deserve further study as possible targets of sexual selection.

## Acknowledgements

The authors extend many thanks to the Tsimane participants and their families, as well as the Tsimane Health and Life History researchers and staff.

## Footnotes

1 Searching “facial width to height ratio” in Google Scholar revealed increasing numbers of publications every year from 2010 (*N*=6) to 2018 (*N*=155).

